# Polarity protein distribution on the metaphase furrow regulates hexagon dominated plasma membrane organization in syncytial *Drosophila* embryos

**DOI:** 10.1101/770453

**Authors:** Bipasha Dey, Debasmita Mitra, Tirthasree Das, Aparna Sherlekar, Ramya Balaji, Richa Rikhy

## Abstract

Epithelial cells have a polarised distribution of protein complexes on the lateral membrane and are present as a polygonal array dominated by hexagons. Metazoan embryogenesis enables the study of temporal formation of the polygonal array and mechanisms that regulate its distribution. The plasma membrane of the syncytial *Drosophila* blastoderm embryo is organized as a polygonal array during cortical division cycles with an apical membrane and lateral furrow in between adjacent nuclei. We find that polygonal plasma membrane organization arises in syncytial division cycle 11 and hexagon dominance occurs with increase in furrow length in cycle 12. This is coincident with DE-cadherin and Bazooka enrichment at edges and the septin, Peanut enrichment at vertices of the base of the furrow. DE-cadherin depletion leads to loss of hexagon dominance. Bazooka and Peanut depletion leads to a delay in occurrence of hexagon dominance from nuclear cycle 12 to 13. Hexagon dominance in Bazooka and Peanut mutants occurs with furrow extension and correlates with increase in DE-cadherin in syncytial cycle 13. We conclude that a change in polarity complex distribution leads to loss of furrow stability thereby changing the polygonal organization of the blastoderm embryo.

**Highlight Summary for TOC:** Metazoan embryogenesis starts with the formation of polygonal epithelial-like cells. We show that hexagon dominance in polygonal epithelial-like plasma membrane organization occurs in nuclear cycle 12 in the syncytial blastoderm *Drosophila* embryo. DE-cadherin and Bazooka distribution along the lateral furrow regulates this hexagon dominance.

## Introduction

Epithelial cells are organised in a polygonal array in various tissues across metazoans. Hexagon dominated polygonal packing is a conserved property of all epithelia across various organisms ranging from the diploblastic *Hydra* to the triploblastic *Xenopus* (Gibson *et al.*, 2006). The epithelial cell plasma membrane (PM) has polarized distribution of protein complexes which allows it to segregate into apical, lateral and basal domains. The Bazooka-Crumbs and Scribble polarity complexes are localized to the apical and basolateral domains, respectively (Laprise and Tepass, 2011). In columnar epithelial cells, the lateral membrane domain may account for up to 60% of the total cell surface area (Tang, 2017). The lateral membrane domains of neighbouring cells adhere to each other with the help of various junctional complexes, thus, contributing to the epithelial cell height. The sub-apical adherens junctions, comprising the E-cadherin plays a significant role in lateral membrane adhesion. Asymmetric distribution of these polarity complexes is important for cell shape, tissue integrity and tissue remodelling (Bilder and Perrimon, 2000; Hayashi and Carthew, 2004; Letizia et al., 2013).

Hexagon dominance, a key feature of several epithelia, has been seen to evolve over developmental stages. For example, in the wing disc of *D. melanogaster*, there is an increase in the percentage of hexagons from 60% to 80% from larval to pupal stages, although the distribution is hexagon dominated from the very beginning (Classen *et al.*, 2005; Sánchez-Gutiérrez *et al.*, 2013). Various molecular and physical factors influence polygonal distribution. Most theoretical models use surface free energy minimization as the constraint that leads to hexagonal packing. Surface energy minimization for a group of epithelial cells is a cumulative result of minimizing the surface area of each cell exposed to the surrounding while maximizing contacts between them, similar to molecules in a fluid bulk (Lecuit and Lenne, 2007). One of the most common molecular factors regulating this distribution is the junctional molecule E-cadherin (E-cad). DE-cad (*Drosophila* E-cad) stabilization due to decrease in turnover in endocytic mutants results in decreasing the frequency of hexagons in *Drosophila* wing discs. This E-cad recycling is shown to be regulated by planar cell polarity (PCP) proteins and the loss of PCP proteins, in turn, shows a decrease in the number of hexagons in the epithelium (Classen *et al.*, 2005; Iyer *et al.*, 2019). In animal cell cultures, it has been observed that loss of ROCK1 and ROCK2, which are important regulators of Myosin II activity, result in shortening of lateral cell height and decrease in the percentage of hexagons (Kalaji *et al.*, 2012). Thus, one of the by-products of polarity might be stability of the lateral membrane and hexagon dominance.

Metazoan embryogenesis shows the onset of epithelial-like polarity and formation of a polygonal array (Nance, 2014). A systematic analysis of onset of polygonal packing and the factors that determine this in embryogenesis has not been characterized thus far. In this study we use the syncytial *Drosophila* blastoderm embryo to characterize the temporal onset of polygonal packing and the role of polarity proteins in regulating its dynamics. Nuclear division cycles (NC) 1-9 occur deep in the interior of the *Drosophila* syncytial blastoderm embryo followed by nuclear migration to cortex during NC10 (Foe and Alberts, 1983; Miller *et al.*, 1985). The arrival of the nuclei can be seen as buds called caps at the cortex (Foe and Alberts, 1983; Miller *et al.*, 1985). In terms of morphological features, the caps are enriched in multiple villi-like projections (Turner and Mahowald, 1976; Miller *et al.*, 1985; Karr and Alberts, 1986; Mavrakis *et al.*, 2009). Nuclear division cycles 11-13 occur beneath the cortex and these projections are reduced at metaphase of each cortical nuclear cycle when the caps are flattened (Turner and Mahowald, 1976). Complete cells that are epithelial in nature are formed in NC14 by extension and polarization of the plasma membrane in a process called cellularization (Lecuit, 2004).

The syncytial *Drosophila* embryo also shows asymmetric distribution of various polarity and cytoskeletal proteins. To begin with, the *Drosophila* embryo is cortically uniform at the pre-blastoderm stage. The first cortical differentiation occurs during NC10 where the cortex is divided into two domains during interphase; the cap and intercap regions. The cap region is enriched in F-actin and actin associated proteins like Arp2/3, SCAR, Moesin, ELMO, Sponge and *α*-spectrin, while the intercap region is marked by Myosin II, Toll and Slam. The PM begins to be organized as a polygonal array and during metaphase, the cortex is further segregated into three domains; apical, lateral and basal domain. In the case of a syncytial cell, a complete basal PM is missing and basal domain refers to the furrow tip. The apical-lateral region is occupied by Canoe, Peanut (Pnut), Scrambled, DE-cad, Bazooka (Baz); lateral by Dlg and Toll; and the furrow tip shows enrichment of PatJ, Amphiphysin, Anilin, Diaphanous and Syndapin (Pesacreta *et al.*, 1989; Thomas and Williams, 1999; Foe *et al.*, 2000; Stevenson *et al.*, 2002; Zallen *et al.*, 2002; Mavrakis *et al.*, 2009; Rikhy *et al.*, 2015; Sherlekar and Rikhy, 2016; Schmidt and Grosshans, 2018; Schmidt *et al.*, 2018).

The syncytial *Drosophila* blastoderm embryo already shows molecular and morphological asymmetries in the PM along with a polygonal array but how these regulate the distribution and dynamics of the polygonal array remain to be investigated. Here, we assess the role of polarity proteins in regulating the polygonal PM organization. We find that the PM of the embryo is organised as a hexagon dominated polygonal array in NC12 with DE-cad and Baz enriched at the edges and Peanut enriched at vertices. DE-cad, Baz and Pnut are enriched at the basal part of the furrow membrane. DE-cad depletion leads to loss of hexagon dominance and a short furrow while Baz and Pnut depletion results in a delay in the onset of hexagon dominance.

## Results

### Onset of hexagon dominated plasma membrane architecture occurs during nuclear cycle 12 in syncytial *Drosophila* blastoderm embryos

Epithelial cells are seen as polygonal cells with pentagons, hexagons and heptagons occuring at high frequencies in various tissues all across the metazoan animal kingdom (Gibson *et al.*, 2006). Hexagon dominance arises as a result of energy minimization while maximizing the area of contact between adjacent cells (Gibson *et al.*, 2006). To assess the time at which this comes about during the syncytial division cycles, we performed live imaging of tGPH expressing embryos. tGPH marks the phospholipid PIP3 enriched PM regions and labelled the PM uniformly in the syncytial *Drosophila* embryo (Britton *et al.*, 2002; Sherlekar and Rikhy, 2016). We used the packing analyzer software to estimate the polygon distribution from metaphase of each NC (Figure 1A).

**Figure 1.**
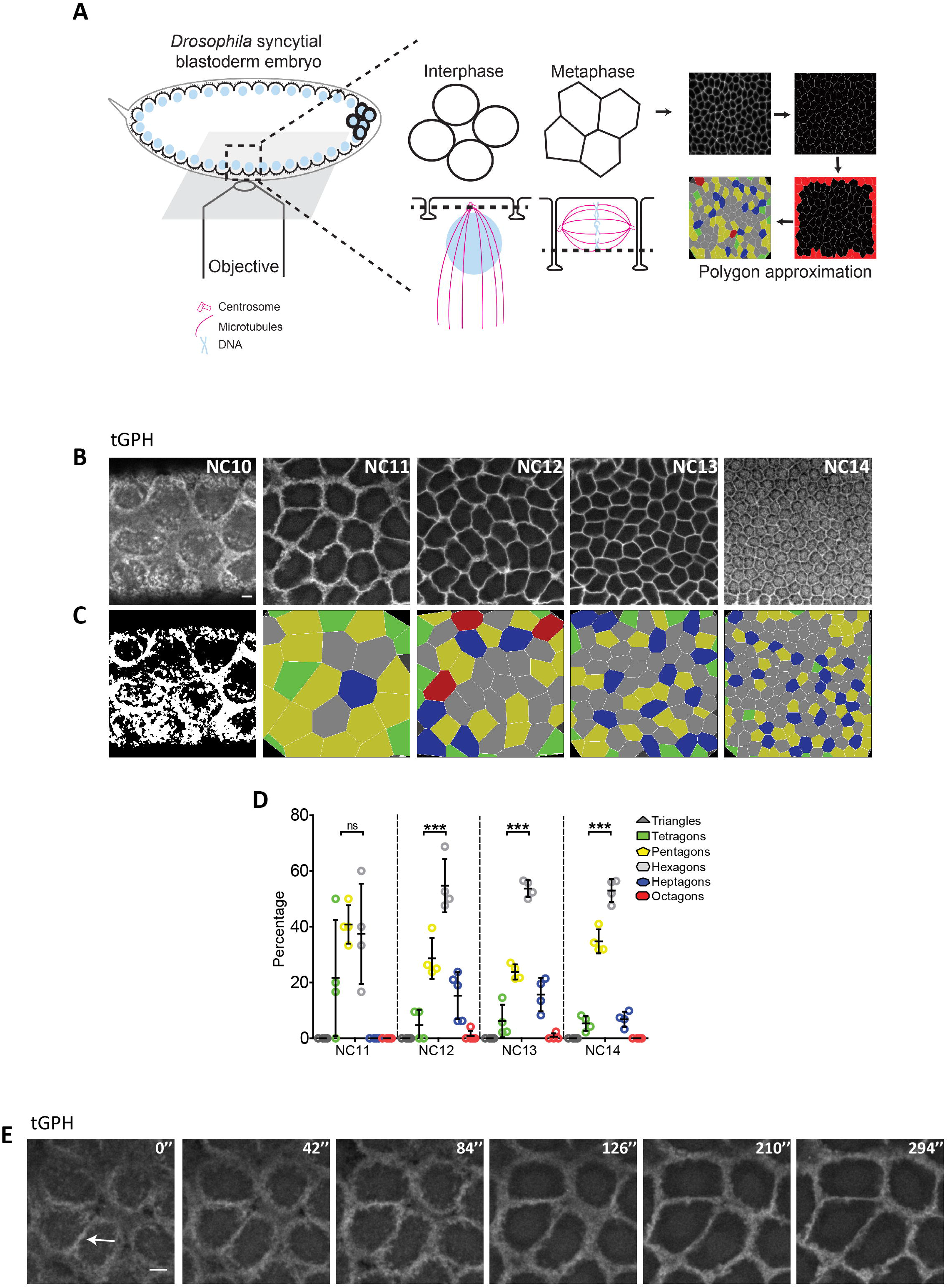
Hexagon dominated plasma membrane organization emerges at NC12. **(A)** Schematic showing the syncytial *Drosophila* blastoderm embryo in interphase being imaged by an objective below in an inverted microscope. The zoomed inset shows a syncytial cell in metaphase turned 180 degrees with the PM on top. A cross-section of this view across the metaphase furrow at the bottom (dotted line) shows polygonal distribution of the cells. These were analyzed by the packing analyzer software for obtaining the polygon distribution onset across the NC10-14. Grazing sections of tGPH expressing embryos from NC10-14 at metaphase, tGPH labels the entire membrane in NC10-13 and enters the nucleus in NC14 in addition to being at the membrane **(B)**. Colour-coded polygon rendering using the packing analyzer software **(C)**. Quantitative analysis of polygon distribution in NC11-14 (n= 20-60 cells from NC11-14 per embryo, 4 embryos) **(D)**. **(E)** Edge formation occurs before vertex formation. Grazing sections of tGPH expressing embryos at NC11 from interphase to metaphase. The arrow shows the formation of edges first, followed by the formation of vertices. Data is represented as mean± SD. NC11 polygon distribution is significantly different from NC12-14 and NC12-14 are similar to each other with hexagon dominance, Multinomial chi square test (***p<0.001). Two tailed, unpaired, Student’s t test is used for comparing hexagons versus pentagons. Scale bar: 5 μm

The nucleo-cytoplasmic domains of the syncytial *Drosophila* embryo have decreased diffusion of organelles and PM proteins across adjacent domains (Frescas *et al.*, 2006; Mavrakis *et al.*, 2009). Hence we refer to these as “syncytial cells”. The syncytial cells were relatively far apart in metaphase of NC10 and seen as separated caps at the embryo surface (Figure 1B, Movie S1). The furrow length increases during each cycle from NC11 to 13 from interphase to metaphase in between adjacent nucleo-cytoplasmic domains and it reaches a maximum at metaphase during syncytial cycle 13 (Foe and Alberts, 1983; Holly *et al.*, 2015; Xie and Todd Blankenship, 2018). The PM was organized into a polygonal array for the first time in *Drosophila* embryo development in NC11 (Figure 1B). The PM in the polygonal array in NC13 and 14 was more taut as compared to NC11 and 12. NC11 at metaphase showed almost equal numbers of pentagons and hexagons in the polygonal array. The polygonal array then became dominated by hexagons followed by pentagons in NC12. This hexagon dominance persisted in NC13-14 (Figure 1C-D). While observing the onset of formation of the polygonal array, we noted that edges formed before vertices in NC11 (Figure 1E).

The furrow length in nuclear division cycles increases from NC11 to 13 (Holly *et al.*, 2015). We found that the furrow length was approximately 7 μm in NC11, 9 μm in NC12 and 11 μm in NC13 with tGPH (Figure 2). Polygonal architecture is also visible at lower furrow lengths in each NC. We chose to analyze the polygon distribution at a lower length of approximately 6.5 μm in NC12 and NC13 when polygonal architecture was clearly visible across the embryo. We found that at this furrow length in NC12, pentagons were dominant, whereas in NC13 hexagons were dominant (Figure S1A-C). These data together show that epithelial-like hexagon dominance first occurs at NC12 at longer furrow lengths in metaphase, even before complete cells are formed in the syncytial division cycles.

**Figure 2.**
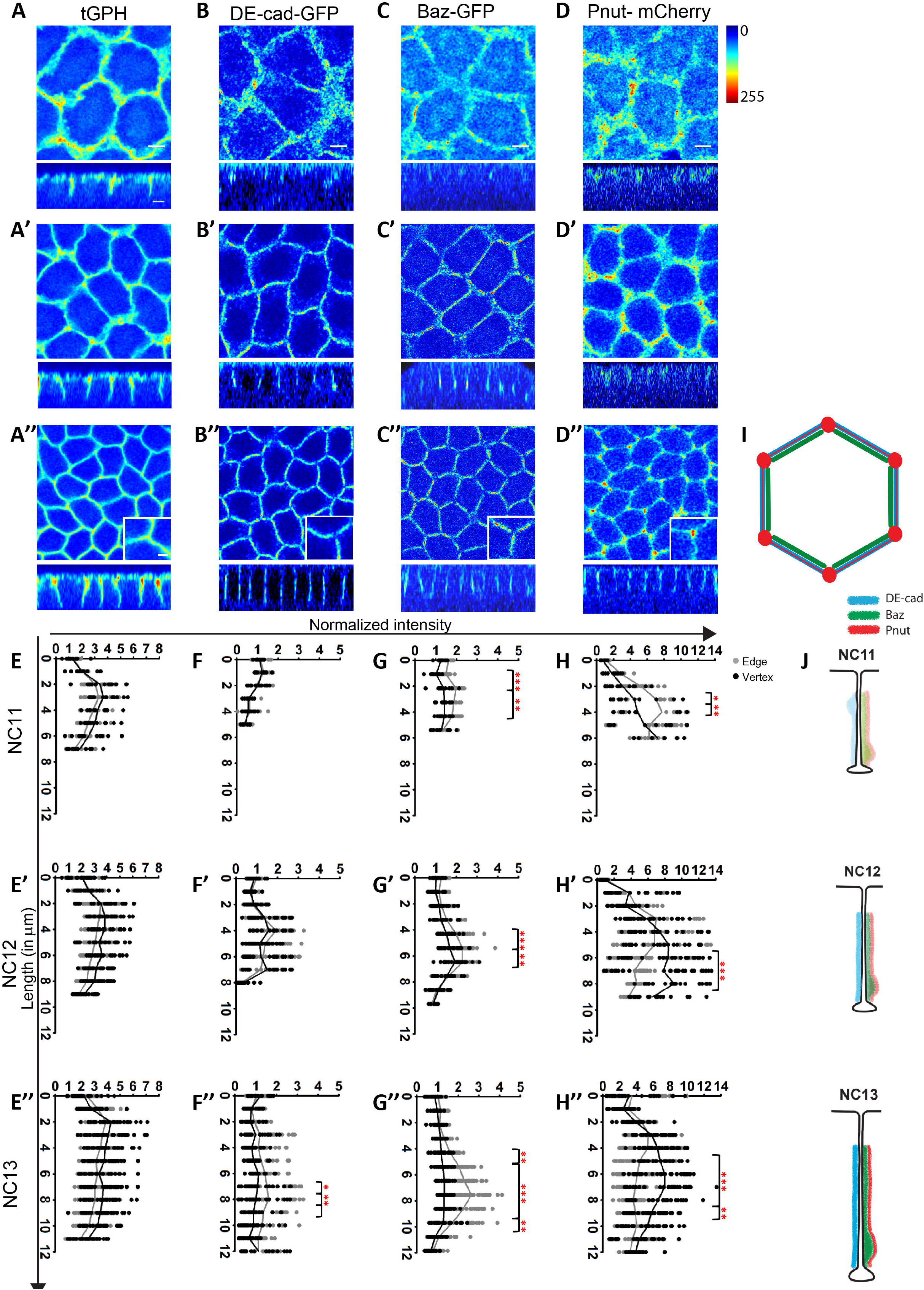
DE-cad, Baz and Pnut show polarized distribution in the syncytial cells. **(A-D)**Distribution of tGPH (**A**: NC11, **A’**: NC12 and **A’’**: NC13), DE-cad-GFP (**B**: NC11, **B’**: NC12 and **B’’**: NC13), Baz-GFP (**C**: NC11, **C’**: NC12 and **C’’**: NC13) and Pnut-mCherry (**D**: NC11, **D’**: NC12 and **D’’**: NC13) in grazing and sagittal views in syncytial NC11-13. The Jet rainbow scale from ImageJ is used to show fluorescent intensities. tGPH labels the entire PM, DE-cad is enriched at edges in NC13 (**B’’**insets show zoomed in images), Baz is enriched at edges NC11 onwards while Pnut is enriched on the vertex NC12 onwards (**C’’-D’’**insets show zoomed in images). **(E-H)**Quantification of intensities normalized to the apical section of NC11 in edges (black) and vertices (grey) along the metaphase furrow length for tGPH (**E**: NC11, **E’**: NC12, **E’’**: NC13), DE-cad-GFP (**F**: NC11, **F’**: NC12, **F’’**: NC13), Baz-GFP (**G**: NC11, **G’**: NC12, **G’**: NC13’) and Pnut-mCherry (**H**: NC11, **H’**: NC12, **H’’**: NC13) in NC11-13 (n=3 embryos each). The total Baz-GFP and tGPH fluorescence on the furrow shows 1.5 fold enrichment on the membrane from NC11-12, the total Pnut-mCherry and DE-cad-GFP fluorescence shows 2 fold enrichment from NC11-12. The total DE-cad-GFP shows 3 fold enrichment on the lateral membrane at NC13 while others do not show further increase. The significance bars along the length of the furrow show significant enrichment on the edge in the basal regions of the furrow for DE-cad-GFP in NC13, for Baz-GFP in NC11-13 and on the vertex for Pnut-mCherry. **(I-J)** Schematic representing the polarized localization of proteins in XZ and XY planes. Asymmetric distribution of DE-cad, Baz and Pnut between edges and vertices in the XY plane **(I)**. Asymmetric distribution of DE-cad, Baz and Pnut across NC11-13 along the XZ plane **(J)**. While DE-cad spreads all across the length, Baz and Pnut are enriched in the basal part of the furrow region. The scatter plots contain a line connecting the means, *p<0.05, **p<0.01, and ***p<0.001, Two way ANOVA with Bonferroni post tests. The stars show significance between edge and vertex intensities at the indicated length based on the post test. Scale Bar=5 μm. The zoomed in insets show a scale bar of 2 μm.

### Analysis of asymmetric distribution of DE-cadherin, Bazooka and Peanut in the plasma membrane of the *Drosophila* syncytial blastoderm

Since the syncytial embryo PM was organized into epithelial-like polygonal array starting from NC11 and became hexagon dominated in NC12, we tested if polarity proteins were progressively enriched in the lateral furrows in NC11-13. DE-cad and Baz have been found in the syncytial furrow and form apical spot junctions in cellularization (Harris and Peifer, 2004). In order to characterize the temporal distribution of DE-cad and Baz as compared to tGPH in NC11-13, we performed live imaging of embryos expressing DE-cad-GFP (Huang et al., 2009) and Baz-GFP (Benton and Johnston, 2003). The epithelial PM shows a distinct distribution of proteins along tricellular junctions, which function in sealing the intercellular space (Ikenouchi et al., 2005; Schulte et al., 2003) as well as along the lateral domain. Also we observed a correlation between hexagon dominance in NC12 with increased furrow length (Figure S1). We therefore quantified the intensity of the fluorescently tagged DE-cad and Baz and the PM marker tGPH in edges and vertices of the polygonal array along with different optical sections in the length of metaphase furrow in NC11-13 (Figure 2A-C). The intensities obtained were plotted as a fold change with respect to the apical section of NC11 (Figure 2E-G). tGPH was distributed evenly across the furrow in edges and vertices and marked the entire length of the furrow in the lateral views (Figure 2A-A’’,E-E’’).

We found that DE-cad was uniform across edges and vertices in NC11 and 12 and was enriched at edges in NC13 (Figure 2B-B’’, F-F”). DE-cad was present along the entire metaphase furrow in NC11-12, while in NC13 it was enriched in the basal part of the furrow from 7-10 μm (Figure 2B-B’’, F-F”).

Baz-GFP was present along the entire membrane with an enrichment at edges in NC12-13. Baz was enriched at the furrow between 2-5 μm in NC11 and towards the basal part of the furrow from 4-6 μm in NC12 and from 6-10 μm in NC13 (Figure 2C-C’’, G-G’’).

The septin family proteins Pnut, Sep1 and Sep2 were studied because Pnut is present at the furrow in syncytial stages and functions in actin organization and furrow extension (Rosalind-Silverman, 2008). Pnut-mCherry (Guillot and Lecuit, 2013) was used to assess the distribution of Pnut on the membrane in edges and vertices and along the length of the furrow across the syncytial cycles. Pnut was present throughout the membrane and was enriched at vertices from NC11 onwards. Pnut was concentrated towards the basal part of the furrow at 3-5 μm in NC11, 5-8 μm in NC12 and 5-10 μm in NC13 (Figure 2D-D’’, H-H”).

We estimated the fold change of total DE-cad, Baz and Pnut fluorescence on the PM across the syncytial cycles as compared to NC11. We found that the total intensity of DE-cad along the metaphase furrow in NC13 as compared to NC12 increased significantly greater than that of Baz and Pnut (Figure 2F” compared to 2G”-H”). This variation in distribution across the syncytial cycles may have implications on the role of these proteins in the formation and stabilization of furrows and polygonal architecture.

We assessed the presence of polarity proteins Crumbs, Stardust, Dlg, Scrib and Patj by immunostaining. Crumbs and Stardust were not expressed in the early embryo (data not shown). Consistent with previous observations, Dlg as present along the furrow membrane while Patj was enriched at the tip (Harris and Peifer, 2004; Mavrakis *et al.*, 2009)(Figure S2A-D). We also found Scrib to be present along the furrow as reported earlier (Schmidt *et al.*, 2018). Edge enrichment was also seen for Dlg and vertex enrichment was observed in Sep1 and Sep2 (Figure S2E-H).

The syncytial PM showed asymmetries in the planar axis of the polygon at edges (DE-cad, Baz) and vertices (Pnut) and enrichment of proteins along the base of lateral furrow from NC11-13 (Figure 2I-J).

### Bazooka membrane recruitment is important for Peanut distribution in the syncytial *Drosophila* embryo

To dissect the role of Baz, Pnut and DE-cad during the onset of hexagon dominance, we investigated their loss of function effects in the syncytial cycles. We assessed the role of Baz by maternally expressing *baz* RNAi (*baz*^i^) (see *material and methods* for details). Baz protein levels as assessed by an antibody against the N-terminus of the protein were lowered in these embryos as compared to control (Wodarz *et al.*, 1999). Interestingly, with the knockdown of this edge enriched protein, Pnut was also lowered (Figure 3A-B) suggesting a possible role of Baz in stabilization of Pnut on the membrane. However, the F-actin remained unaltered and appeared similar to control embryos (Figure 3A’-B’). As Baz function is important for formation of spot adherens junctions in cellularization (Harris and Peifer, 2004), we checked DE-cad distribution in *baz*^i^ embryos and found that its distribution was similar to controls in NC13 (Figure 3B’).

**Figure 3.**
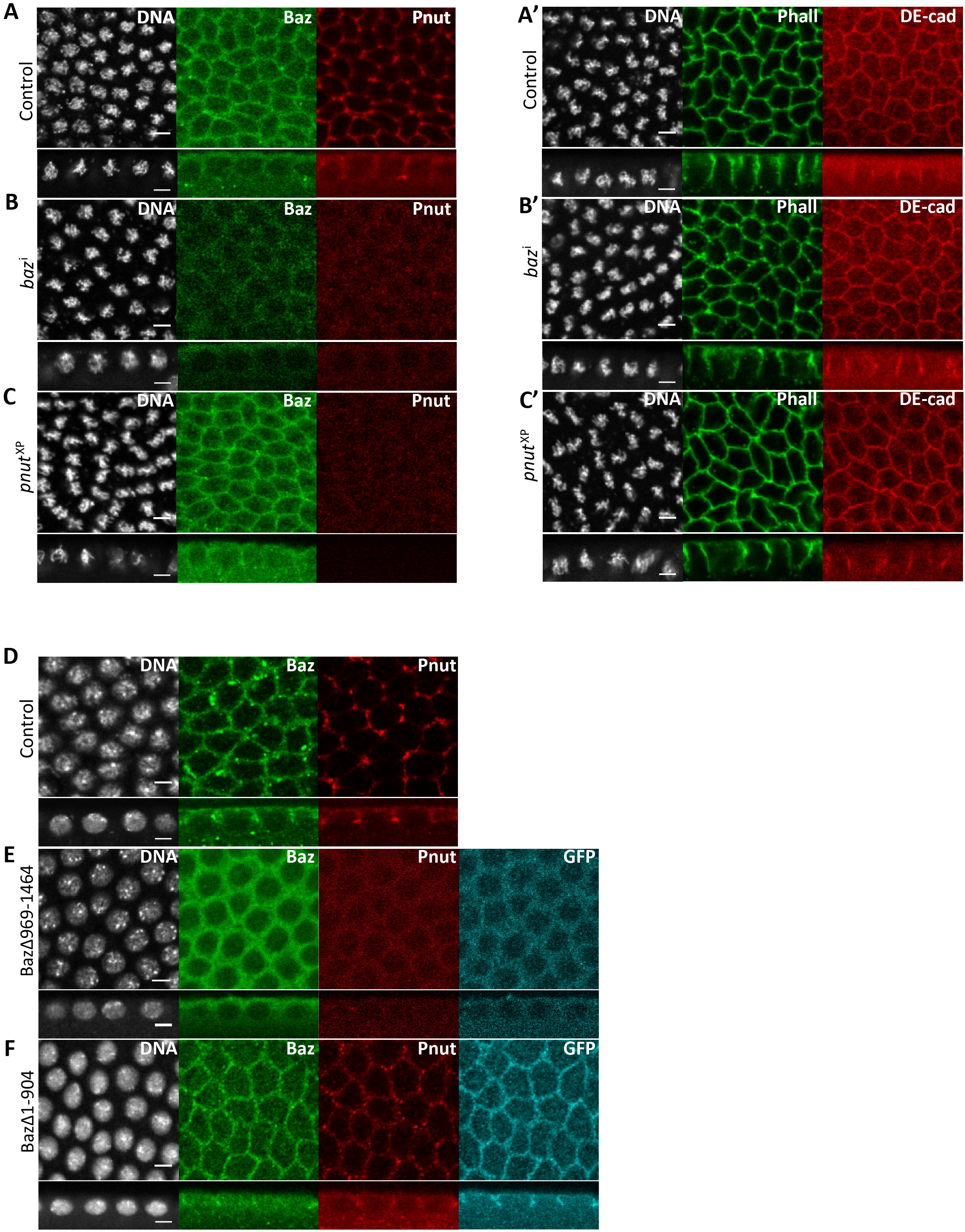
Baz membrane binding domain is important for Pnut recruitment on the membrane. **(A-C, A’-C’)** *mat*-Gal4/+ **(A)** control (n=20), *baz*^i^ **(B)**and *pnut*^XP^ **(C)** embryos co-stained with Baz (green) and Pnut (red) **(A-C)**; DE-cad (red) and phalloidin (green) **(A’-C’)**. *baz*^i^ shows a decrease in Baz and Pnut (100%, n=20) **(B)**, DE-cad is unaffected (100%, n=10) **(B’)**. *pnut*^XP^ shows loss of Pnut (100%, n=17) **(C)**; Baz (100%, n=11) and DE-cad (100%, n=15) are unaffected **(C-C’)**. **(D-F)** Overexpression of Baz truncated for the PM binding domain decreases Pnut recruitment to the membrane. Control (n=10 embryos) **(D)**, Baz∆969-1464-GFP **(E)** and Baz∆1-904-GFP **(F)** expressing embryos co-stained with Baz (green), Pnut (red) and GFP (cyan). Baz∆969-1464-GFP shows cytosolic distribution and Baz antibody shows membrane and cytosolic localization **(E)**. These embryos also show loss of Pnut from the membrane (87%, n=24 embryos). Baz∆1-904-GFP and Baz antibody shows membrane localization **(F)**. Pnut distribution is slightly reduced but vertex enrichment is present (64%, n=14 embryos). DNA (grey). Scale Bar=5 μm

To determine the effect of loss of Pnut on Baz and DE-cad localization, we generated germline clone embryos of null mutants of Pnut (*pnut*^XP^) (Neufeld and Rubin, 1994). With depletion of Pnut from *pnut*^XP^ embryos, Baz and DE-cad localization remained unaffected (Figure 3A,C,A’,C’). Thus, Pnut was not important for Baz localization on the syncytial PM. Notably, Baz and Pnut depletion did not affect DE-cad distribution on the syncytial PM.

To verify if Pnut localization on the syncytial PM depended on Baz association to the syncytial PM, we maternally overexpressed truncated transgenes of Baz containing the N terminus oligomerization domain or the C terminus phospholipid membrane binding domain. A GFP tagged C-terminal truncation mutant of Baz, BazΔ969-1464-GFP, which is defective in PM recruitment (Krahn et al., 2010) was expressed maternally in the presence of wild-type protein. BazΔ969-1464-GFP showed a cytosolic pattern in metaphase of NC13. Baz antibody staining against the N-terminus of the protein showed an increased cytosolic distribution and a diffused membrane localization as compared to controls (Figure 3D-E). Since Baz is seen to form oligomers in *vivo* (Benton and St Johnston, 2003), we speculate that the N-terminal domain oligomerizes in this overexpression mutant, leading to the reduction of Baz from PM and an increase in the cytosol in addition to the attenuated levels of Pnut. On the other hand, when a GFP tagged Baz N-terminal truncation mutant, BazΔ1-904-GFP was maternally overexpressed, both the truncated and endogenous Baz could localize on the membrane. Pnut distribution was weaker than that seen in controls but it was present on the membrane (Figure 3D,F). Thus, the C-terminal domain of Baz was important not only for its own recruitment on the PM but also for Pnut membrane localization.

### Bazooka and Peanut depletion leads to delayed onset of hexagon dominance and short furrows in the syncytial blastoderm embryo

We analyzed polygonal distribution in Baz and Pnut knockdown embryos, by live imaging mutant embryos expressing tGPH and with phalloidin staining. We found that *baz*^i^ and *pnut*^XP^ (Figure 4A-B,E) showed a significant increase in the frequency of pentagons and loss of hexagon dominance in NC12. However, the hexagon dominance was seen similar to controls in NC13 (Figure 4C-D,E). Overexpression of the Baz oligomerization domain (BazΔ969-1464-GFP) that lowered Baz and depleted Pnut from the membrane also showed loss of hexagon dominance at NC12 which recovered at NC13 (Figure S3A-D).

**Figure 4.**
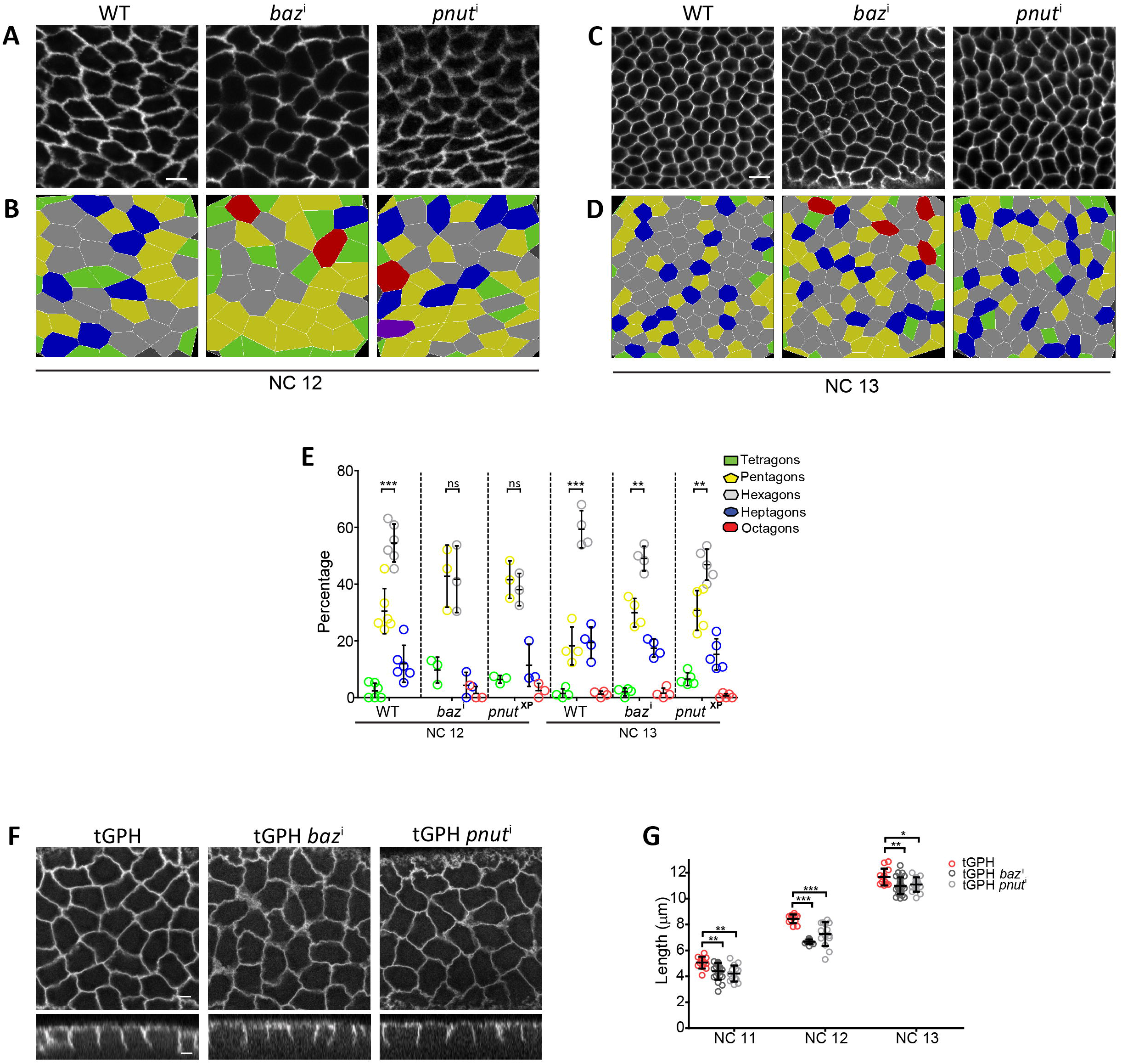
Baz and Pnut depletion show delay in hexagon dominance and decreased furrow length. **(A-D)** Wild-type, *baz*^i^ and *pnut*^XP^ embryos stained with phalloidin in NC12-13 **(A,C)** along with the respective colour-coded polygon renderings **(B,D)**. **(E)** Polygon distribution in the mutants and wild-type stained with phalloidin in NC12-13. Polygon distributions of wild-type and *baz*^i^ and *pnut*^XP^ embryos are significantly different from each other (*p<0.05) at NC12 but not at NC13. Multinomial chi square test (n=approx. 60-80 syncytial cells, 20-30 cells/embryo; 4-5 embryos). Hexagon dominance in *baz*^i^ and *pnut*^XP^ recovers in NC13. Pentagons and hexagons are compared in each cycle using the two tailed, unpaired, Student’s t test. Data is represented as mean ± SD, *p<0.05, **p<0.01, and ***p<0.001. **(F-H)** *baz* and *pnut* mutant embryos show decreased furrow length. tGPH grazing sections in control, *baz*^i^, *pnut*^i^ and *baz*^i^ *pnut*^i^ at NC12 **(F)**. Quantification of metaphase furrow lengths in tGPH/+, *baz*^i^ and *pnut*^i^ in NC11-13 (n=12, 4 furrows; 3 embryos) **(G)**. Data is represented as mean ± SD, *p<0.05, **p<0.01, and ***p<0.001, Two tailed, unpaired Student’s t test. Scale bar: 5 μm.

Since we found increased localization of Baz and Pnut at the base of the metaphase furrow, we estimated the furrow length in *baz*^i^ and *pnut*^i^ embryos. Pnut loss is known to result in shorter metaphase furrows (Rosalind-Silverman, 2008; Sherlekar and Rikhy, 2016). *baz*^i^ and *pnut*^i^ embryos showed a marginal but significant decrease in furrow lengths in NC11-13 as compared to controls (Figure 4F-G, Movie S2-S3). We also estimated the furrow ingression rates of the knockdowns and observed similar rates of ingression in both mutants as compared to the control even though the final length was slightly decreased (Figure S3E). The double mutant of Baz and Pnut showed a loss of furrow length like the single mutant and this was also decreased to a significant but small extent (Figure S3F-G). Thus, Baz and Pnut mutant embryos showed a marginal decrease in furrow lengths metaphase of the NC11-13 and a delay in onset of hexagon dominance.

### DE-cadherin depletion leads to loss of hexagon dominance and severely disrupted furrow extension

We assessed DE-cad mutant embryos for localization of Baz and Pnut along with polygon onset in the syncytial cycles. *shg* RNAi (*shg*^i^) was maternally expressed in embryos (see Materials and Methods for details). As expected, embryos developing to syncytial stages had reduced DE-cad staining (Figure 5A-B). The *shg*^i^ embryos had diffuse F-actin distribution as seen by phalloidin staining when compared to sharp staining in controls. Baz and Pnut were present on the furrow membrane in *shg*^i^. However, Pnut was spread more evenly and not enriched at the vertex and Baz staining was more diffuse as compared to controls (Figure 5A’-B’). DE-cad distribution of the membrane was therefore important for Baz and Pnut localization on the furrow membrane.

**Figure 5.**
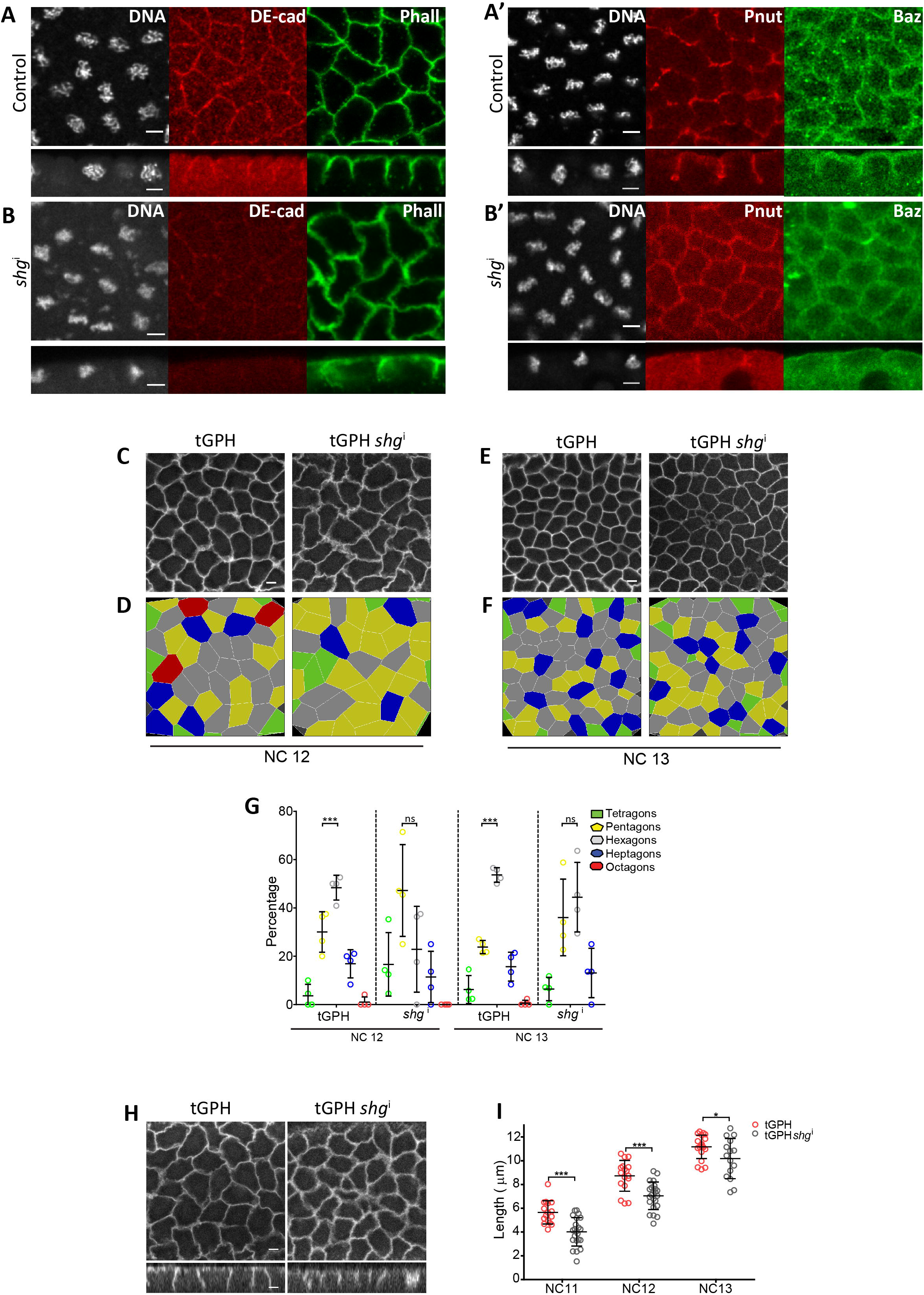
DE-cad depletion results in mislocalization of Baz and Pnut, loss of hexagon dominance and decreased furrow length. **(A-B, A’-B’)** DE-cad is lowered in *shg*^i^ embryos. *nanos*-Gal4/+ (n=10) and *shg*^i^ (**B**, 86%, n=21) embryos are stained with DE-cad (red) and phalloidin (green) **(A-B)**; Baz (green) and Pnut (red) **(A’-B’)**. Phalloidin, Baz and Pnut are more spread in *shg*^i^ embryos (94%, n=16) and the sharp distribution is lost. **(C-G)***shg*^i^ embryos show loss of hexagon dominance. tGPH/+ and tGPH *shg*^i^ embryos in NC12-13 **(C,E)** along with the respective colour-coded polygon renderings **(D,F)**. Graph showing polygonal distribution in *shg*^i^ in NC12-13 **(G)**. The polygon distributions of *shg*^i^ are significantly different from control (*p<0.05) using Chi square test (n=approx. 120 syncytial cells, 20-30 cells/embryo; 4 embryos). Hexagon dominance is not seen in NC13. Pentagons and hexagons are compared in each cycle using the unpaired, two tailed, Student’s t test. The NC13 polygon distribution for the control is repeated from Figure 1D. **(H-I)** *shg*^i^ embryos show decreased furrow length. tGPH grazing sections from control (n=4 embryos) and *shg*^i^ with short furrow lengths at NC12 (61%, n=18 embryos) **(H)**. Graph showing quantification of metaphase furrow lengths in tGPH/+ and *shg*^i^ embryos in NC11-13 (n=12, 4 furrows; 3 embryos) **(I)**. Data is represented as mean ± SD, *p<0.05, **p<0.01, and ***p<0.001, Two tailed, unpaired Student’s t test. Scale Bar = 5 μm.

To assess the effect of DE-cad loss on polygonal shape, we quantified the polygonal distribution from metaphase of *shg*^i^ live movies obtained with tGPH. Live imaging of *shg*^i^ embryos with tGPH showed defects during syncytial division cycles. *shg*^i^ embryos showed loss of hexagon dominance at NC12. *shg*^i^ expressing embryos did not show a significant difference between pentagons and hexagons in NC13. Thus, unlike *baz*^i^ and *pnut*^i^ embryos, loss of hexagon dominance persisted in NC13 in *shg*^i^ (Figure 5C-G). *shg*^i^ expressing embryos also showed ruffled membranes as opposed to the taut and sharp membranes in the controls. The furrow membrane showed a diffuse tGPH signal which was spread over a larger area as compared to control embryos (Figure S3H-I, Movie S4). Therefore, DE-cad loss affected polygon distribution more than Baz and Pnut.

Finally, we estimated furrow length in *shg*^i^ embryos with tGPH and found it to be considerably shorter than the control and shorter on average than *baz* and *pnut* knockdowns (Figure 5H-I compared to 4F-G). Taken together, DE-cad plays a significant role in maintaining hexagon dominance. This occurs by its known function of mediating adhesion between adjacent PM lateral domains and in the case of the syncytial embryo, adjacent furrow membranes together for the formation of edges.

### Recovery of hexagon dominance in Bazooka and Peanut mutant embryos occurs during syncytial cycle 13 with furrow extension

Furrows increase in length from interphase to metaphase in each NC (Xie and Todd Blankenship, 2018). Baz and Pnut depletion resulted in delayed hexagon dominance and marginally short furrows. DE-cad depletion, on the other hand, had loss of hexagon dominance and gave shorter furrows than Baz and Pnut loss. Since Baz and Pnut mutant embryos showed a recovery of hexagon dominance in NC13, we estimated whether hexagon dominance occurred as a function of furrow length in NC13. We plotted the distribution of polygons and ratio of pentagons to hexagons in controls, Baz and Pnut mutant embryos with respect to furrow length. For control embryos, we found that hexagon dominance was present as soon as the polygonal array was established in NC13 at 6.5 μm and remained the same until metaphase (Figure 6A-F). Interestingly, Baz and Pnut mutants showed pentagon dominance at the start of NC13 in interphase and showed hexagon dominance at the maximum furrow length in NC13 (Figure 6A-F). In summary, the recovery to hexagon dominance in NC13 in both Baz and Pnut knockdown embryos was furrow length dependent. It is possible that increased recruitment of polarity complexes occurred at increased furrow length and this allowed the membranes to stabilize to give rise to hexagon dominance.

**Figure 6.**
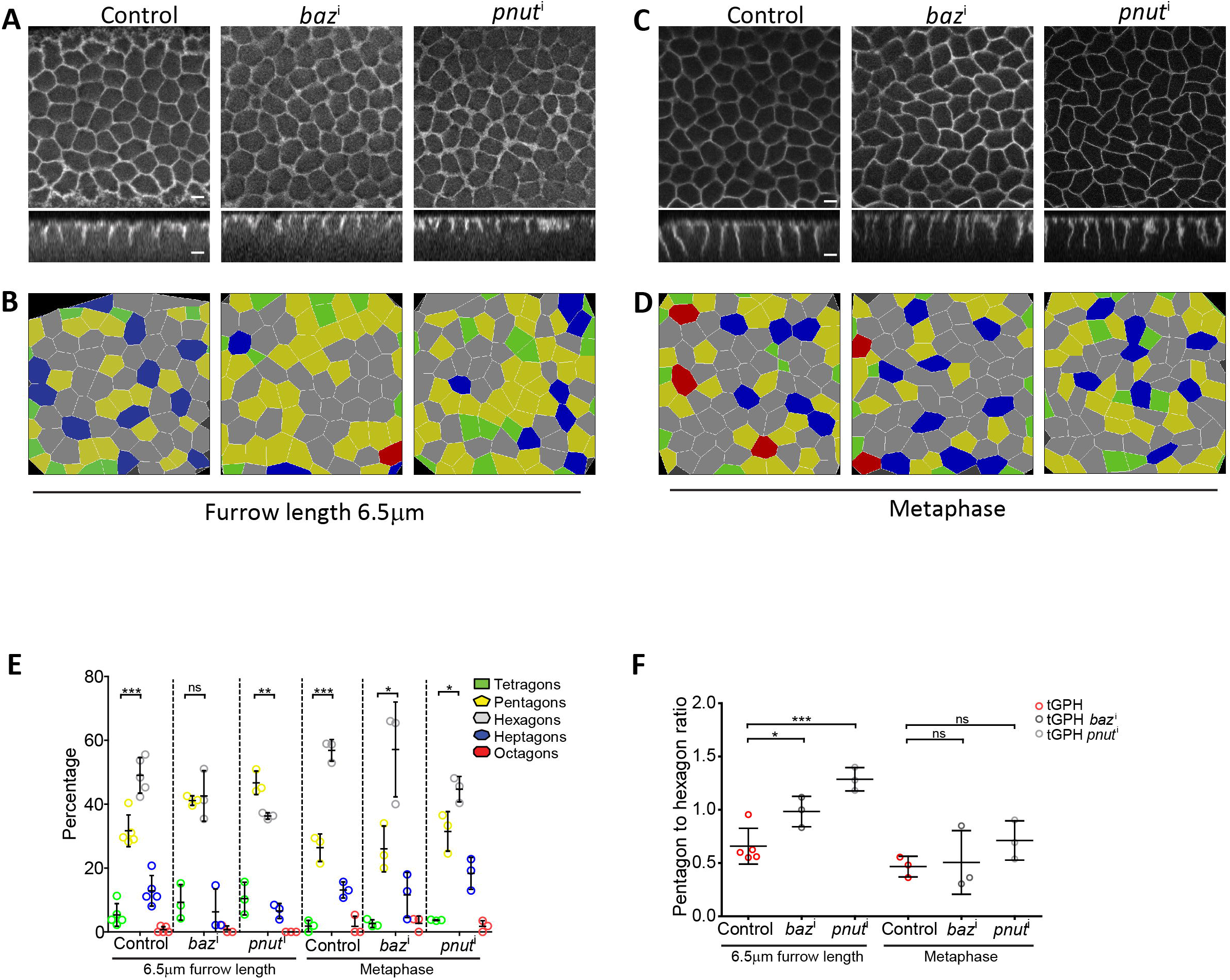
Appearance of hexagon dominance in Baz and Pnut knockdowns occurs at an increased furrow length in NC13. **(A-B)** *baz*^i^ and *pnut*^i^ embryos show loss of hexagon dominance at a shorter furrow length at NC13 when wildtype is already hexagon dominant. Grazing and sagittal sections of tGPH expressing control, *baz*^i^ and *pnut*^i^ embryos at NC13 at a short furrow length of 6.5 μm **(A)** along with the respective colour-coded polygon renderings **(B)**. **(C-D)** *baz*^i^ and *pnut*^i^ embryos show recovery of hexagon dominance at the maximum furrow length at metaphase NC13. Grazing and sagittal sections of tGPH expressing control, *baz*^i^ and *pnut*^i^ embryos at NC13 at the maximum furrow length at metaphase **(C)** along with the respective colour-coded polygon renderings **(D)**. **(E-F)** Baz and Pnut knockdowns show a length dependent recovery of hexagon dominance at NC13. Graph showing the polygon distribution of control, *baz*^i^ and *pnut*^i^ embryos at NC13 at a short furrow length of 6.5μm and at metaphase **(E)**. Graph showing the pentagon to hexagon ratio of control, *baz*^i^ and *pnut*^i^ embryos at NC13 at a short furrow length of 6.5 μm and at metaphase **(F)**. Data is represented as mean ± SD, *p<0.05, **p<0.01, and ***p<0.001, Two tailed, unpaired Student’s t test. Scale Bar=5 μm

### Occurence of hexagon dominance in Bazooka and Peanut mutant embryos in NC13 is coincident with increase in DE-cadherin

Apical cap remodelling in the syncytial division cycles occurs with the help of regulators of actin remodelling. The actin caps are formed in interphase of syncytial division cycles and expand to form furrows in prophase and metaphase. Arp2/3 activity is needed for cap expansion followed by Myosin II which leads to cap buckling for the formation of furrows (Stevenson *et al.*, 2002; Zhang *et al.*, 2018). Also Anillin-Pnut networks have been found to play a redundant function to Myosin II in furrow initiation (Zhang *et al.*, 2018). Septins have also been found crucial for bundling actin into curved bundles in cellularization (Mavrakis *et al.*, 2014). Our data shows that furrow formation is affected in embryos depleted of DE-cad possibly due to lack of adhesion and stabilization of furrows. We find that loss of DE-cad leads to increase in pentagons. Thus, it is possible that change in levels of Myosin II and/or DE-cad levels at the furrow leads to recovery of hexagon dominance in Baz and Pnut mutant embryos.

We hence imaged Myosin II and DE-cad dynamics in Baz, Pnut and DE-cad depleted embryos. For imaging Myosin II we expressed the fluorescently tagged Myosin light chain subunit Spaghetti Squash (Sqh) tagged with either GFP or mCherry (Royou *et al.*, 2004; Martin *et al.*, 2009). As reported previously, we found that Myosin II was enriched on the PM in interphase and was depleted from the PM in metaphase (Figure 7A-C). DE-cad was present on the membrane in both interphase and metaphase of NC11 and 12. Increased activation of Myosin II on increasing RhoGEF2 activity leads to increased recruitment of fluorescently tagged Sqh on the membrane (Izquierdo *et al.*, 2018). We analyzed the Myosin II and DE-cad levels by imaging Sqh-mCherry; DE-cad-GFP in Baz and Pnut depleted embryos and Myosin II levels by imaging Sqh-GFP in DE-cad depleted embryos. We estimated the total Sqh-mCherry or Sqh-GFP fluorescence on the furrow as a ratio to the neighboring cytoplasm. We did not find a significant difference in interphase in Baz, Pnut and DE-cad depleted embryos as compared to controls (Figure 7D-E,D’-E’,G-H). Sqh was decreased from the membrane in Baz, Pnut and DE-cad depleted embryos in metaphase similar to controls (data not shown).

**Figure 7.**
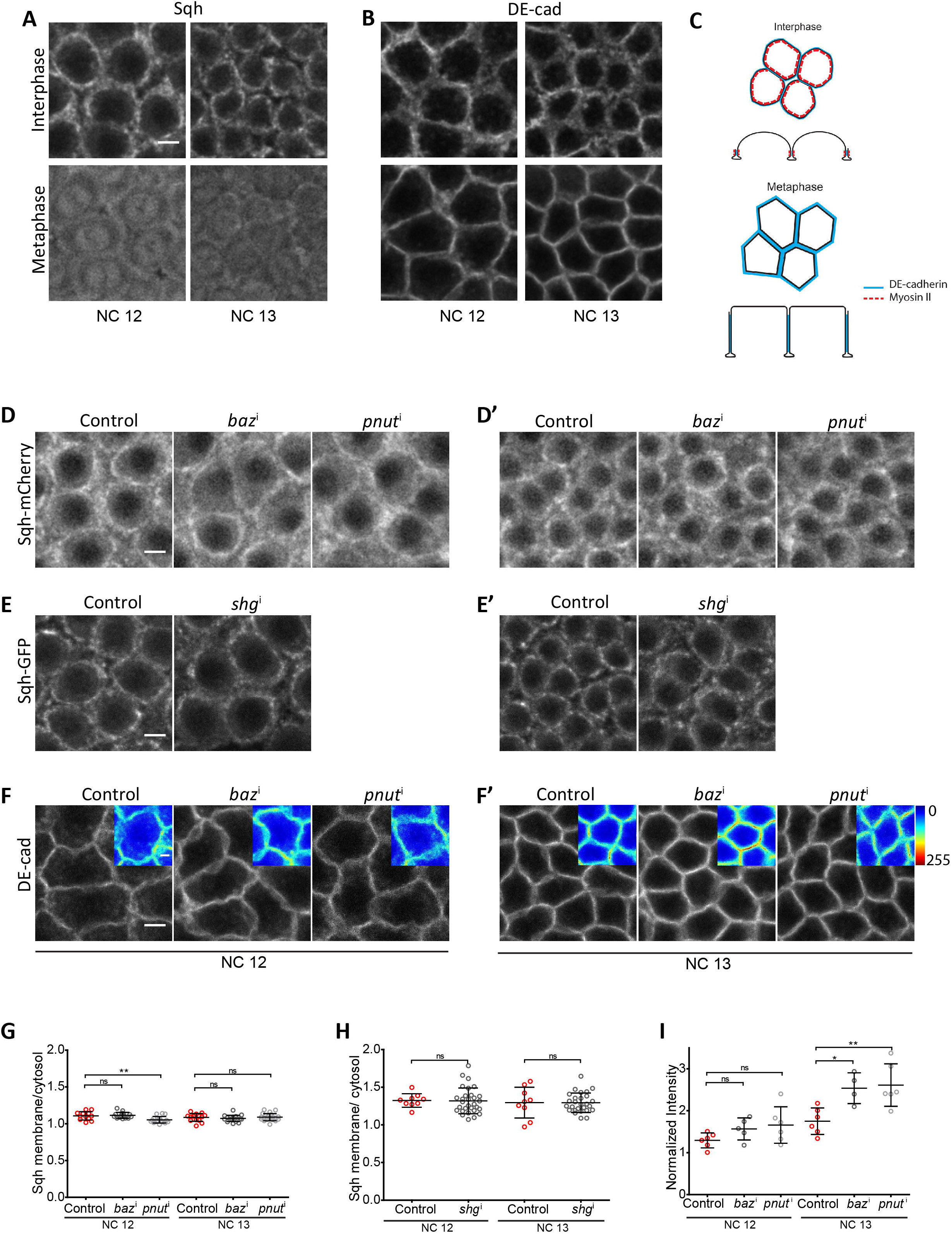
Appearance of hexagon dominance in Baz and Pnut knockdowns correlates with increased levels of DE-cad in NC13. **(A-C)** Myosin II and DE-cad distribution in NC13 in control embryos. Grazing sections of Sqh-mCherry **(A)** and DE-cad-GFP **(B)** expressing embryos at interphase and metaphase of NC12-13. Schematic representing the MyoII and DE-cad localization. DE-cad is on the membrane in interphase and metaphase while MyoII becomes cytosolic in metaphase **(C)**. **(D-I)** MyoII levels remain unchanged and DE-cad levels increase in NC13 in Baz and Pnut knockdowns. Sqh-mCherry and DE-cad-GFP coexpressing control and Baz and Pnut depleted embryos were used for quantification of Sqh and DE-cad in NC12-13. Sqh-GFP was expressed in DE-cad depleted embryos for quantification of Sqh in NC12-13. Grazing section of Sqh-mCherry expressing control, *baz*^i^ and *pnut*^i^ embryos at interphase in NC12-13 **(D,D’)**. Grazing sections of Sqh-GFP expressing control and *shg*^i^ embryos at interphase in NC12-13 **(E,E’)**. Note that Sqh-mCherry is generally more cytoplasmic as compared to Sqh-GFP. Hence the membrane to cytoplasm ratios for Sqh-mCherry were lower Sqh-GFP. Grazing sections of DE-cad-GFP expressing control, *baz*^i^ and *pnut*^i^ embryos at metaphase in NC12-13 **(F,F’)**. Graph comparing MyoII membrane to cytosol ratio between control, *baz*^i^ and *pnut*^i^ in NC12-13 (n=12-15, 5 cells per embryo, 3 embryos)**(G)**. Graph comparing MyoII membrane to cytosol ratio between control and *shg*^i^ in NC12-13 (n=12-15, 5 cells per embryo, 3 embryos) **(H)**. Graph comparing DE-cad fold change with respect to NC12 between control, *baz*^i^ and *pnut*^i^ (n=12-15, 5 cells per embryo, 5 embryos) **(I).** Data is represented as mean ± SD, *p<0.05, **p<0.01, and ***p<0.001, Two tailed, unpaired Student’s t test. Scale Bar=5 μm. The zoomed-in insets show scale bar of 3 μm.

DE-cad levels were next estimated in Baz and Pnut mutant embryos by expressing DE-cad-GFP in these mutant embryos. We found that Baz and Pnut embryos did not show a significant change in DE-cad levels as compared to controls in NC12. We next represented the total DE-cad fluorescence in NC13 as a ratio to NC12 and found that there was a distinct increase in DE-cad levels in Baz and Pnut depleted embryos as compared to controls (Figure 7F,F’,I).

In summary, we observe that loss of DE-cad leads of loss of hexagon dominance (Figure 5) and increase of DE-cad in Baz and Pnut depleted embryos correlates with occurence of hexagon dominance during NC13 (Figure 4,7).

## Discussion

In this study we show that hexagon dominated plasma membrane organization occurs in the syncytial *Drosophila* blastoderm embryo from NC12 onwards. Pentagons and hexagons are equally likely in NC11 when edges first form and hexagons are present at almost double the number of pentagons from NC12 onwards. Since the syncytial cycles do not have a complete basal domain the mechanisms that regulate this hexagon dominance are likely to be present on the lateral furrow. We have characterized the role of Baz, Pnut and DE-cad proteins in regulation of the lateral furrow length in the syncytial embryo. This analysis reveals furrow and polygon distribution phenotypes in two categories: 1) DE-cad depleted embryos have short furrows and loss of hexagon dominance; 2) Baz and Pnut depleted embryos have only slightly short furrows and delay in hexagon dominance. Whereas in the control embryos, hexagon dominance appears in NC12 at longer furrow lengths, hexagon dominance appears in Baz and Pnut mutant embryos in NC13 with increase in furrow lengths and coincident with an increase in DE-cad levels. These studies therefore reveal the combination of polarity proteins necessary to give rise to a *de novo* hexagon dominant epithelial-like PM in the syncytial *Drosophila* embryo (Figure 8).

**Figure 8.**
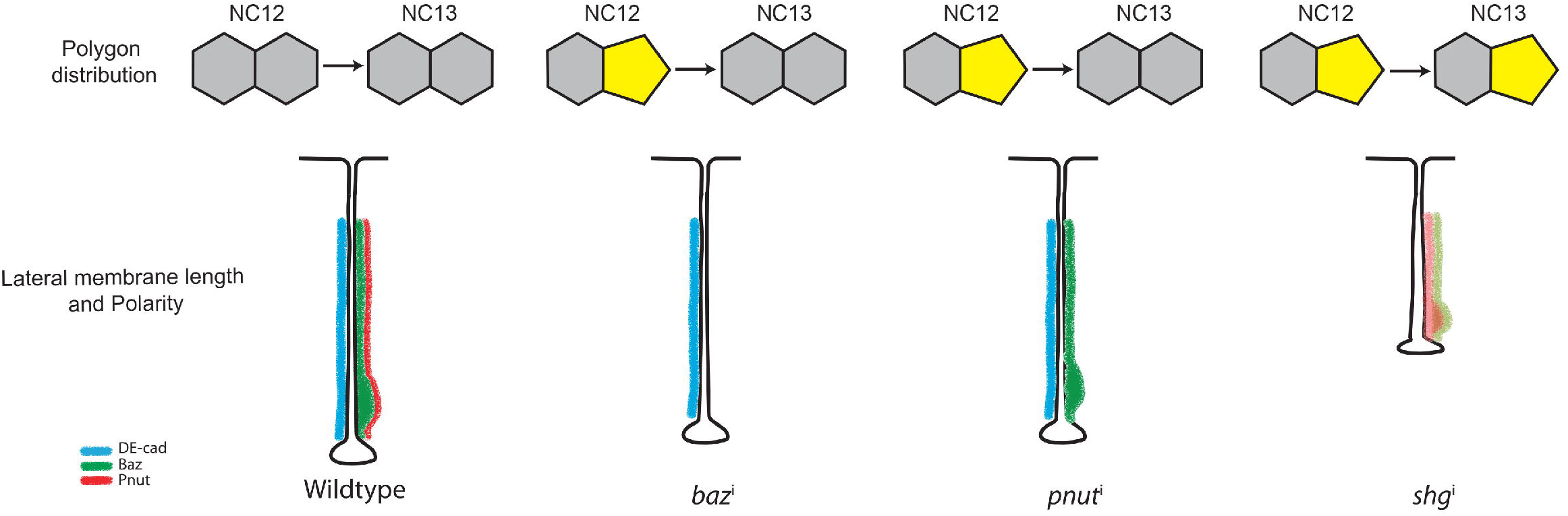
Summary schematic showing the distribution of DE-cad, Baz and Pnut on the lateral furrow and polygon distribution in the *Drosophila* syncytial blastoderm embryo. Control embryos show hexagon dominance in NC12-13 with significant amount of DE-cad, Baz and Pnut on the lateral furrow. Loss of Baz or Pnut results in a delay in the onset of hexagon dominance while loss of DE-cad results in loss of hexagon dominance. Note the difference in the protein complex composition of the various mutants. *baz*^i^ shows loss of Pnut in addition to loss of Baz, while *pnut*^i^ shows loss of Pnut only. DE-cad is unperturbed in both these cases. It instead shows an increase at NC13 which correlates with the recovery of hexagon dominance. *shg*^i^, on the other hand, shows lowering and mislocalization of Baz and Pnut.

DE-cad loss decreased but did not completely abolish lateral furrows. If this is the major protein responsible for stabilization of adhesion of furrow membranes of adjacent syncytial cells, we should have obtained a phenotype of complete loss of furrow but we only saw a severe reduction in furrow length. We argue that this could be due to inability to deplete DE-cad completely with the RNAi strategy. Also it could be due to the presence of other proteins that are responsible for keeping the furrow membrane adhered to each other. We tested for the occurence of other transmembrane proteins such as Crumbs, Neuroglian and Neurexin (Harris and Peifer, 2004; Laprise *et al.*, 2009) and did not find them to be present on the furrow in syncytial embryos. Future analysis of more such cadherin-like adhesion molecules such as Echinoid may be useful in this direction (Wei *et al.*, 2005).

Increase in DE-cad is known to occur due to the loss of endocytosis and recycling in the *Drosophila* wing disc epithelium (Classen *et al.*, 2005; Iyer *et al.*, 2019). Dynamin dependent endocytosis also regulates the syncytial furrow dynamics and DE-cad on the furrow membrane (Rikhy *et al.*, 2015). It is possible that loss of Baz and Pnut on the syncytial furrow leads to increase in DE-cad on the furrow in NC13 due to loss of recycling. Further analysis on changes in membrane trafficking on the PM on the loss of a polarity proteins will enable ascertaining decreased endocytosis in Baz and Pnut depleted embryos as a mechanism for reversal of hexagon dominance in Baz depleted embryos.

Baz and DE-cad play significant roles in initiating the polarity program in different tissues. Baz initiates adherens junction polarity in *Drosophila* cellularization and gastrulation (Müller and Wieschaus,1996; Harris and Peifer, 2004; (Pilot *et al.*, 2006)) but is dispensable in follicle epithelial cells (Shahab et al., 2015). Conversely, in mesoderm invagination, apical movement of DE-cad precedes Baz relocation and thus, the asymmetry in DE-cad distribution here does not depend on Baz (Weng and Wieschaus, 2017). This is similar to mammalian cells where E-cad is recruited to contact points before Baz (Coopman and Dijane, 2016). This suggests that depending on the tissue type or developmental stage, the relative importance of Baz and E-cad in initiating polarity may change. We show that DE-cad is important for distribution of Baz and Pnut to the furrow. As mentioned above, we were unable to identify the presence of other transmembrane junctional proteins, and in such a scenario, DE-cad is likely to assume a significant role in furrow formation in the *Drosophila* syncytial blastoderm embryo.

Asynchronous cell division in *Drosophila* wing disc epithelia is one of the mechanisms that gives rise to a hexagon dominated and energy minimized network (Gibson et al., 2006). It is of interest to note that synchronous division in the syncytial *Drosophila* blastoderm embryo also reaches a similar hexagon dominance in NC12. Decreased number of edges in the polygonal array is a favorable state for cell neighbor exchanges and gives rise to a soft network (Farhadifar et al., 2007). The increase in pentagons in *baz* and *pnut* mutants indicates a similar transition to a soft network possibly due to decreased stabilization of the furrow even in the presence of DE-cad and furrow length. The contacts in the Baz and Pnut mutant embryos are likely to facilitate neighbor exchanges. With increase in DE-cad on the furrow in NC13, these may allow for conversion back to hexagon dominated state by formation and stabilization of one additional edge and vertex. Thus, future analysis of change in furrow tension in various genetic backgrounds along with mathematical modelling will reveal the mechanisms that drive shape morphogenesis in *Drosophila* syncytial blastoderm embryo.

## Abbreviations used

PM: plasma membrane
NC: nuclear cycle
DE-cad: DE-cadherin
MyoII: Myosin II
*shg*: shotgun
*baz*: bazooka
*pnut*: peanut
Sqh: Spaghetti Squash

## Acknowledgements

We thank Mandar Inamdar (IIT, Bombay, India) for discussions on this work. We thank the RR lab members for comments and discussion on the data in the manuscript. We thank the *Drosophila* and Microscopy facilities at IISER, Pune, India for help with stocks and microscopy for the experiments. We thank Benoit Aigouy for sharing the Packing analyzer software. We thank Andreas Wodarz for Bazooka antibody and Bazooka domain truncation stocks. We thank Manos Mavrakis for *peanut* mutant stocks and Sep1 and 2 antibodies. BD, DM, and AS thank CSIR for graduate fellowship. RR thanks IISER, Pune, DBT and DST for funding to the lab. RB and TD thank KVPY for fellowship.

## Materials and Methods

### Fly stocks, crosses and lethality estimation

*Drosophila melanogaster* stocks were raised in standard cornmeal agar at 25 °C and 29 °C for RNAi experiments. Embryos obtained from CantonS flies or CantonS flies crossed to *maternal* ⍰*-tubulin* Gal4-VP16 (*mat*-Gal4) or *nanos*-Gal4 (*nos*-Gal4) were used as control. Maternal driver line *mat67;mat15* carrying *maternal* ◻*4 tubulin-*Gal4-*VP16* (obtained from Girish Ratnaparkhi, IISER, Pune, India), homozygous for chromosome II and III was used for all RNAi and overexpression experiments except for *shg*^i^. Baz RNAi (Bloomington Stock number #35002), Pnut RNAi (#65157), DE-cad RNAi (#38207), tGPH (#8163), UASp-Baz-GFP (#65845) and endo-DE-cad-GFP (#60584) lines were obtained from the Bloomington Stock Center, Indiana, Bloomington, USA. ubi-cad-GFP was obtained from the Maithreyi Narasimha lab from TIFR, Mumbai, India. *pnut*^XP^ FRTG13/CyO and *UASp*-Pnut-mCherry stocks were obtained from Manos Mavrakis, Fresnel University, Marseilles, France. Baz truncation domain constructs were obtained from Andreas Wodarz, Goettingen University, Germany. *Sqh*-Sqh Cherry, *mat67*-Gal4; *Ubi*-DE-cad-GFP, *mat15*-Gal4/TM3Sb was obtained from Adam Martin’s lab, MIT, Massachusettes, USA. *Sqh*-Sqh GFP, *mat67*-Gal4 from Thomas Lecuit, IBDM, Marseilles, France.

F1 flies were put in a cage for egg collection to perform immunostaining or live imaging. Germline clones of *pnut*^XP^ were made by crossing *ovo*^D^ FRTG13 males to *hsflp; GlaBlc* females to obtain hsflp; *ovo*^D^ FRTG13/*GlaBlc* males.These males were then crossed to *pnut*^XP^ FRTG13/*Cyo* females. Larvae, pupae and adults emerging from this cross were heat shocked at 37.5 °C. hsflp*;ovo*^D^ FRTG13/*pnut*^XP^ FRTG13 adults were then put in a cage to collect embryos depleted of *pnut*. *shg*^i^ was crossed to a single chromosomal copy of *nos*-Gal4 and maintained at 18 °C to lower the severity of phenotype and obtain fertilized eggs to perform experiments. F1 flies expressing *shg*^i^ with *nos*-Gal4 when grown at 25 or 29 °C, laid embryos that were arrested early in the pre-blastoderm stage of development and, hence, the experiments were performed at 18 °C to allow for Gal4 dilution. This cross at 18 °C gave enough embryos that entered the syncytial cycles. The lethality of *shg*^i^ embryos was 100% (n=150) at 25 °C and 29 °C and 70% (n=200) at 18 °C after 24 hours. The lethality of *pnut*^i^ and *baz*^i^ expressing embryos and *pnut*^XP^ germline clones was 100% (n=300 embryos each).

### Immunostaining

0-2.5 hr old embryos were collected on sucrose agar plates, washed and dechorionated with 100% bleach for 1 min. Embryos were then fixed using 1:1 mixture of 4% Paraformaldehyde in PBS and Heptane for 20 min. Fixed embryos were then either hand-de-vitellinized for phalloidin staining or MeOH de-vitellinized, washed thrice in 1X PBST (1X PBS with 0.3% Triton X100) and blocked in 2% BSA (Sigma-Aldrich, India) in 1X PBST for 1 hr. Primary antibody was then added at an appropriate dilution and incubated overnight, followed by three 1X PBST washes, and 1hr incubation in appropriate fluorescently coupled secondary antibodies at 1:1000 (Molecular probes, Bangalore, India). Hoechst 33258 was added for 10 min in 1X PBST. Finally, the embryos were washed three times in 1X PBST and mounted in Slow fade Gold antifade reagent (Molecular Probes). The primary antibodies used were: rabbit anti-Baz (1:1000 from Andreas Wodarz, Germany), mouse anti-Pnut (1:5, DSHB), mouse anti-Dlg (1:100 DSHB), rabbit anti-Patj (1:1000 from Hugo Bellen, USA), rat Sep1 (1:250 from Manos Mavrakis, France), guinea pig Sep2 (1:250 from Manos Mavrakis, France), rat anti-DE-cad (1:5, DSHB), DNA was stained with Hoechst 33258 (1:1000, Molecular Probes, Bangalore, India).

### Live Imaging of *Drosophila* embryos

1-1.5 hr old embryos expressing the membrane marker tGPH or Sqh-GFP or Sqh-mCherry; DE-cad-GFP were collected and dechorionated with 100% bleach for 1 min and mounted on coverslip-bottomed LabTek chambers (Mavrakis *et al.*, 2008). The chambers were then filled with 1X PBS and imaged on Zeiss Plan Apochromat 40X/ 1.4 NA oil objective.

### Microscopy

Live or fixed embryos were imaged on any of the following laser scanning confocal microscopes: Zeiss LSM710, LSM780 and Leica SP8. The 40X objective with NA 1.4 was used to image living and fixed embryos. The Argon laser was used to image GFP in tGPH, DE-cad, Baz-GFP and Sqh-GFP. The Diode 561 laser was used to image the Sqh-mCherry and Pnut-mCherry. Care was taken to maintain the laser power and gain with the range indicator mode such that the 8-bit image acquired did not show any saturation and was within the 0-255 range. Averaging of 2 was used for both fixed and live imaging. Images were acquired with an optical section of 1.08 microns in all except actin stainings where an optical section of 0.34 microns was used.

### Embryo Lethality

3-4hr old embryos were collected, washed and arranged into a 10 × 10 matrix on a sugar-agar plate using a brush. The number of unhatched embryos were counted after 24 hrs. This procedure was repeated 3 times for each genotype tested.

### Quantification and statistical analysis

#### Image quantification

##### Polygon analysis

The most taut and bright grazing section from metaphase (usually at the base of the furrow) of each NC per embryo was used to quantify polygons using the Packing analyzer software (Benoit Aigouy, Classen et al., 2005, https://idisk-srv1.mpi-cbg.de/~eaton/). The software allows to outline each cell and colour code it according to polygon type. It also provides an excel sheet with the area, perimeter and polygon type of each cell in the field. 3 or more embryos were used per NC and all the cells in the field were analysed this way to obtain the polygon distribution per cycle.

##### Quantification of relative fluorescent signal across depth and in planar sections of the plasma membrane

The grazing sections at metaphase across depth expressing various polarity proteins were used for this analysis. ROIs were drawn at 5 edges and 5 vertices for NC11, and 10 edges and 10 vertices for NC12-13, in each optical section from apical to basal sections. Optical sections were taken at approximately 1 μm depth across the entire furrow. The mean intensities from these ROIs were measured using ImageJ. The intensities from each stack were background subtracted. The graphs shown in Figure 2 represent mean intensities obtained across the depth of the furrow normalized to the mean intensity of the apical-most optical section in NC11.

For calculating relative fold change of fluorescence for DE-cad-GFP, Baz-GFP and Pnut-mCherry in the furrow in NC11 to NC13, total intensities across the furrow length were computed by summation of the mean intensity across the number of optical stacks in NC11, 12 and 13. The total intensities obtained for the entire furrow in NC12 and 13 were divided by the total intensity obtained in NC11 to obtain fold change.

Quantification for Dlg, Sep1 and Sep2 was done by creating 5 ROIs on the edge and the vertex as mentioned above. The mean intensities were normalized by dividing by the mean intensity of the entire field.

Enrichment in the basal furrow region and edge/vertex were computed by performing statistical analysis for the fluorescence values on the length of the furrow using two way ANOVA to test how the intensity of signals varied with either edge/vertex or apical/basal regions of the furrow.

##### Quantification of the metaphase furrow length

Metaphase furrow lengths were quantified from the orthogonal sections for different time points and NCs using the Zen blue software. Approximately 5-8 furrows were measured per time point per embryo. These lengths were further confirmed with the number of z stacks taken to cover the entire furrow length of syncytial cells in the field of view.

### Statistical analysis

All data are represented as mean ± SD. Statistical significance was determined using the Two-tailed, unpaired, Student’s t-test in most cases, to compare between two means. One way ANOVA was used when comparing three or more means together, with Dunnett’s Multiple Comparison Test as post test to compare all means to a control. Multinomial Chi square test was used to compare the polygon distributions between control and mutants in addition to the Student’s t test to compare hexagons versus pentagons for checking hexagon dominance.

## Data and software availability

All raw data in the form of movies and images is available on request from the corresponding author.

