## Supplementary legends for "Polarity protein distribution on the metaphase furrow regulates hexagon dominated plasma membrane organization in syncytial *Drosophila* embryos"

**Affiliation and contact information**

Biology, Indian Institute of Science Education and Research, Homi Bhabha Road, Pashan, Pune, 411008, India Phone: +91-20-25908065

Present address:

1: Present address: Biology, University of Iowa, 246 Biology Building, 129 E. Jefferson St. Iowa City, Iowa 52242-1324, USA

2: Present address: Department of Biology, Stanford University, Stanford, CA, USA

3: Present address: Albert-Ludwigs-University Freiburg, Center for Biological Systems Analysis, Habsburgerstrasse 49, 79104 Freiburg, Germany

**Figure S1**

**Hexagon dominance occurs in NC12 at a longer furrow length.**

**(A-C)** Grazing sections of tGPH expressing embryos from NC11-13 at furrow length just above the threshold (approximately 6.5µm) **(A)** and the respective colour-coded polygon rendering using the packing analyzer software **(B)**. Quantitative analysis of polygon distribution in NC11-14 (n=4 embryos) **(C)**. NC14 polygon approximation was done manually due to weak signal on the membrane and to avoid errors in segmentation. The same control picture and data of N13 are also used to compare to Baz and Pnut mutant embryos in Figure 6A. Frequencies of hexagons and pentagons are significantly different at NC13-14. Data is represented as mean ± SD, *p<0.05, **p<0.01, and ***p<0.001, Two tailed, unpaired, Student’s t test. Scale Bar = 5 µm.

**Figure S2**

**Asymmetric localization of polarity regulators and septin family proteins in the syncytial embryo .**

**(A-D)** Grazing and sagittal sections of wild-type embryos stained with Dlg **(A),** Scrib **(B) a**nd PatJ **(C)** at NC13. Graph showing edge enrichment of Dlg **(D)**. Note the sagittal section in Dlg staining shows enrichment in the basal portion of the furrow while Patj shows basal tip enrichment.

**(E-H)**The septin family proteins Sep1 and Sep2 show vertex enrichment in the syncytial plasma membrane in NC13-14. Grazing sections of embryos expressing Sep2-GFP along with the antibody staining **(E)** and Sep1-GFP along with the antibody staining **(F)** in wild-type embryos at NC13 **(A)**. Graphs show the comparison of signal on edge versus vertex in Sep2-GFP **(G)** and Sep1-GFP **(H)** expression embryos at single optical section (n=15, 5 edges and vertices from each embryo; 3 embryos). Data is represented as mean ± SD, *p<0.05, **p<0.01, and ***p<0.001, Two tailed, unpaired, Student’s t test. Scale Bar = 5 µm.

**Figure S3**

**Characterization of polygon distribution in Baz truncation mutant and membrane architecture in DE-cad knockdown embryos**

**(A-D)** Polygon distribution shows a delay in hexagon dominance in Baz mutant embryos. Grazing sections of Baz∆969-1464-GFP mutant embryos stained with phalloidin at NC12 **(A)** and NC13 **(C)** and the respective colour-coded polygon renderings as obtained from packing analyzer. Comparison of the polygonal distribution in the mutants with wild-type phalloidin stained images at NC12 **(B)** and NC13 **(D)**. Data is represented as mean ± SD, *p<0.05, **p<0.01, and ***p<0.001. The polygon distributions of wild-type, *baz*^i^, Baz∆969-1464-GFP and *pnut*^XP^ mutant embryos are significantly different from each other (*p<0.05) at NC12 but not at NC13 (n=approx. 60 syncytial cells, 20-30 syncytial cells from each embryo; 3 embryos) as tested with Multinomial Chi square test. Hexagons and pentagons are compared using Two tailed, unpaired Student’s t test. The same graph as Figure 4E is used to compare the Baz∆969-1464-GFP truncation mutant.

**(E-G)** Rate of furrow ingression does not change in *baz*^i^ and *pnut*^i^ embryos **(E)**. The furrow length was estimated with *baz*^i^, *pnut*^i^ double mutant embryos generated with *mat*-Gal4 expressing tGPH **(F)**. Quantification of metaphase furrow lengths in *baz*^i^ *pnut*^i^ double knockdown and tGPH/+ in NC12-13 (n=12, 4 furrows; 3 embryos) **(G).**

Data is represented as mean ± SD, *p<0.05, **p<0.01, and ***p<0.001.

**(H-I)** *shg*^i^  embryos show loose and ruffled membrane in syncytial cells. Grazing sections of tGPH expressing *shg*^i^ embryo **(H)**. Line profiles of (yellow line) control and *shg*^i^ embryos. DE-cad loss shows a broader peak width **(I)** at NC12 as compared with controls, which suggests loose and ruffled membranes.

Scale Bar = 5 µm.

**Supplementary Movies**

**Movie S1**

**tGPH imaging across syncytial cycles in apical and orthogonal planes**

The movie shows plasma membrane dynamics in living embryos expressing tGPH during the syncytial division cycles. The planar X-Y view across the syncytial cells is in the center while the top and right panels show the orthogonal views along Y-Z and X-Z, respectively. At NC10 there is enrichment of tGPH signal in apical caps on the plasma membrane above the nuclei. The apical caps are separated from each other across the embryo. Note the onset of polygonal organization in the plasma membrane during the syncytial cycles beginning from NC11. Scale Bar = 5 µm.

**Movie S2**

***baz*^i^ embryos have a shorter metaphase furrow**

The syncytial cycles were imaged in *baz* RNAi expressing embryos with tGPH. The furrow lengths are shorter than the tGPH/+ control. Scale Bar = 5 µm.

**Movie S3**

***pnut*^i^ embryos have a shorter metaphase furrow**

The syncytial cycles were imaged in *pnut* RNAi expressing embryos with tGPH. The furrow lengths are slightly shorter than the tGPH/+ control. Scale Bar = 5 µm.

**Movie S4**

***shg*^i^ expressing embryos have short furrows and membrane ruffling.**

The syncytial cycles were imaged in *shg* RNAi expressing embryos with tGPH. The furrow lengths are much shorter than the tGPH/+ control and NC11-12 show loose and ruffled membrane. Scale Bar = 5 µm.

**Movie S5**

**DE-cad GFP and Sqh-mCherry expressing embryos.**

The syncytial cycles were imaged in embryos expressing DE-cad GFP and Sqh-mCherry. Note cyclical recruitment of Sqh on the membrane at interphase and fall off at metaphase while DE-cad is retained on the membrane throughout.

**Movie S6**

***baz*^i^ embryos expressing DE-cad GFP and Sqh-mCherry**

The syncytial cycles were imaged in *baz* RNAi expressing embryos with DE-cad GFP and Sqh-mCherry.

**Movie S7**

***pnut*^i^ embryos expressing DE-cad GFP and Sqh-mCherry**

The syncytial cycles were imaged in *pnut* RNAi expressing embryos with DE-cad GFP and Sqh-mCherry

**Movie S8**

***shg*^i^ embryos expressing Sqh-GFP**

The syncytial cycles were imaged in *shg* RNAi expressing embryos with Sqh-GFP.
