## Supplementary figures and images for "Polarity protein distribution on the metaphase furrow regulates hexagon dominated plasma membrane organization in syncytial *Drosophila* embryos"

### Figure S1

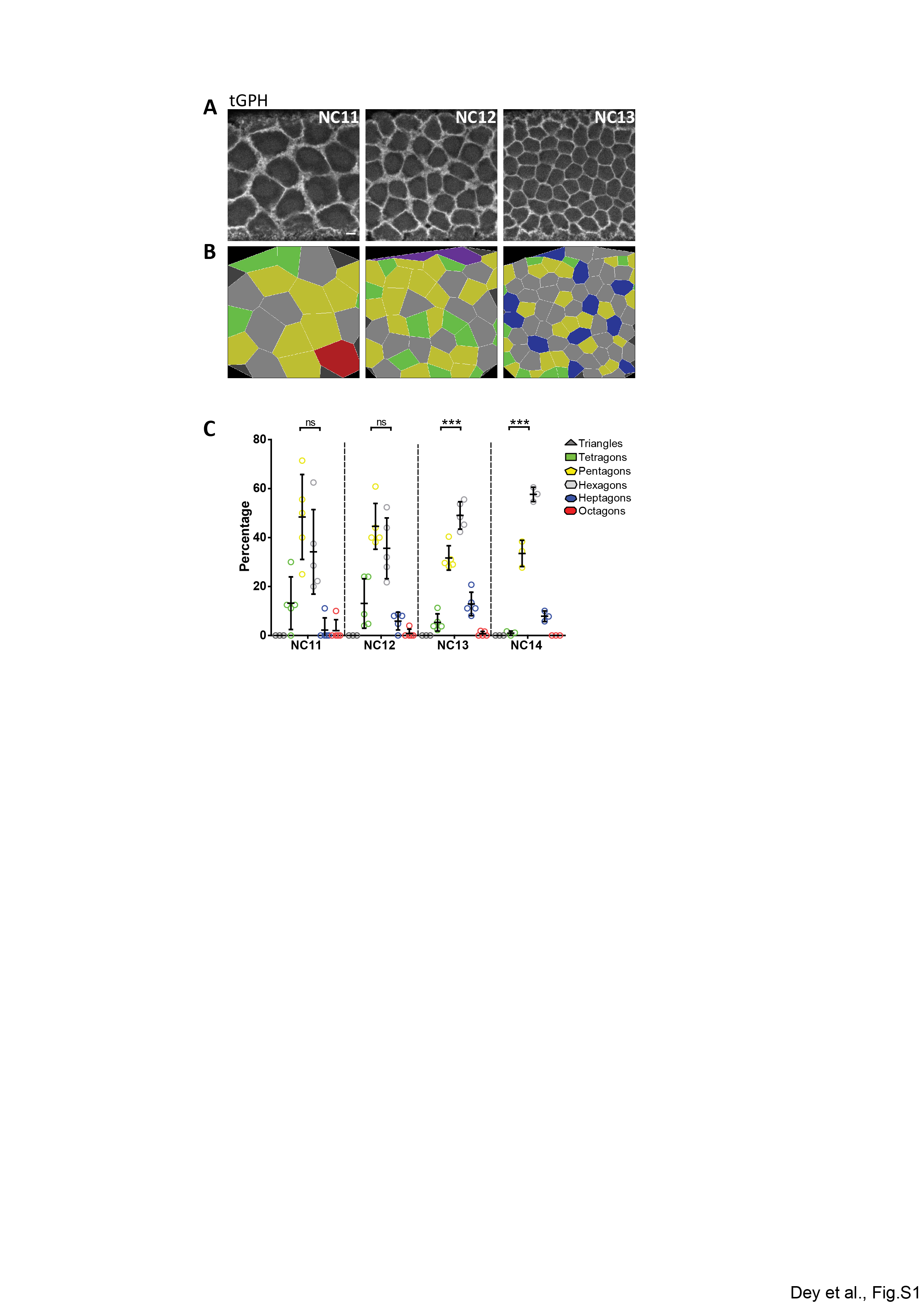

### Figure S2

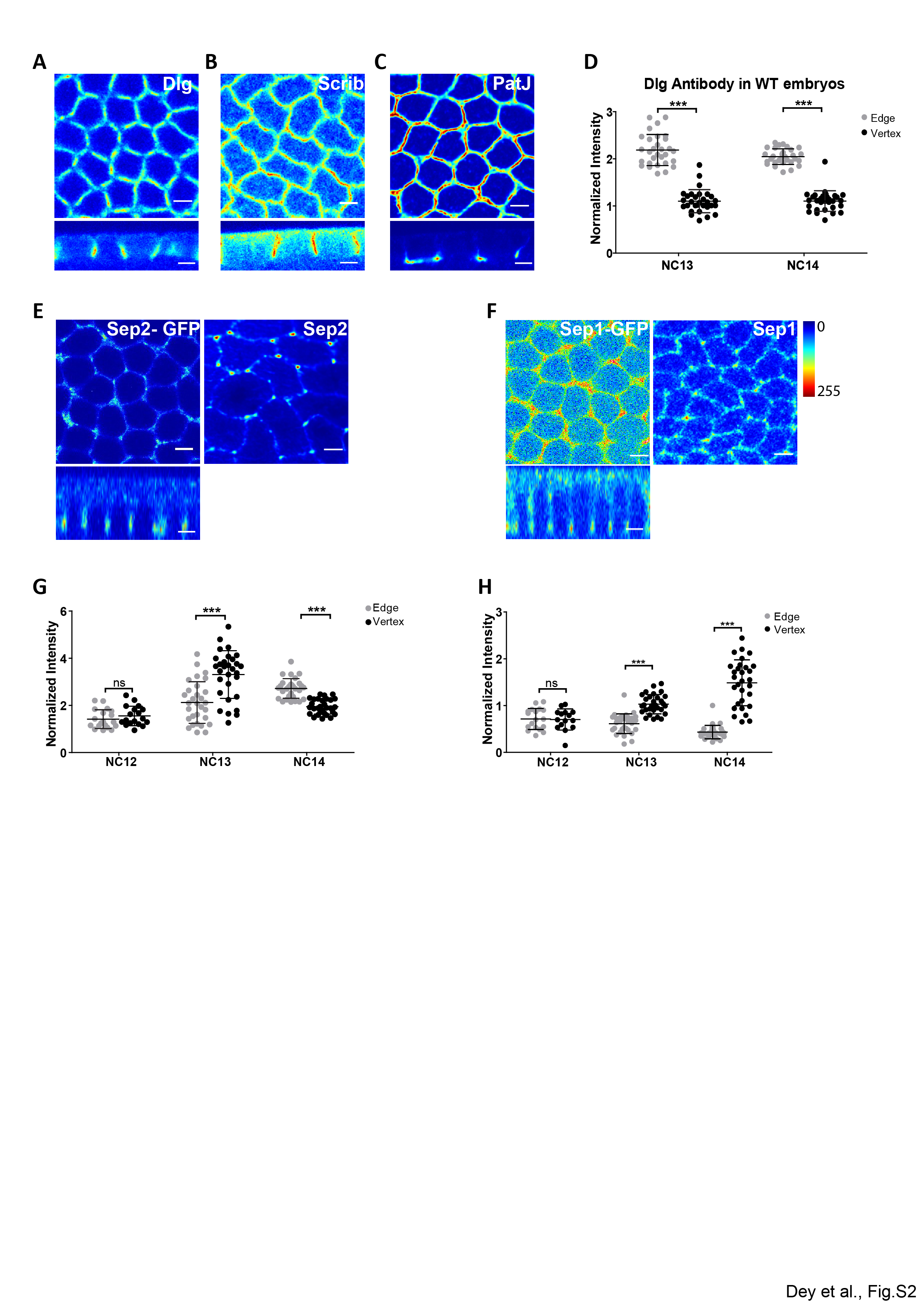

### Figure S3

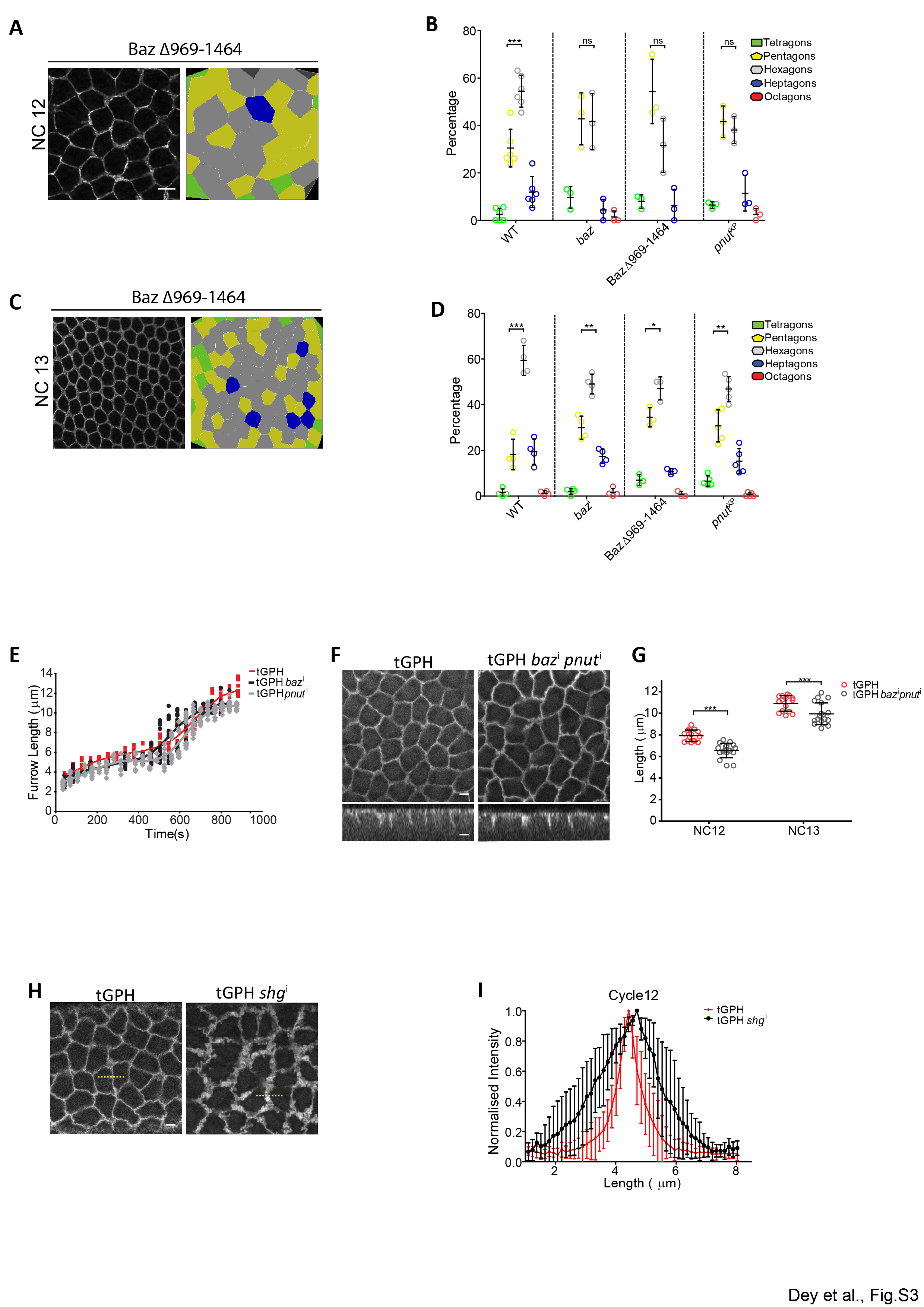
